# Systems analysis reveals alternate metabolic states adopted by *Mycobacterium tuberculosis* across species

**DOI:** 10.1101/2025.11.15.688116

**Authors:** Jonathan Padilla-Gomez, Rachel McGinn, Yisha Liang, Kendra Libby, Alison E. Ringel, Charles Evavold, Bryan D. Bryson

## Abstract

*Mycobacterium tuberculosis* (Mtb) persists within macrophages, yet how different host species shape bacterial state remains poorly understood. Here, we directly compared the intracellular transcriptome of Mtb during infection of human and mouse macrophages, revealing distinct host-imposed microenvironments that drive the pathogen into separable metabolic states. Lipid metabolism and regulatory circuits were prominently remodeled, with mouse macrophages inducing iron- and oxidative-stress responses while human macrophages promoted fatty acid import programs. Using fluorescent fatty acid tracing, we uncovered a striking species-specific phenotype: Mtb forms intracellular lipid inclusions (ILIs) in murine macrophages but not in human macrophages. This phenotype was independent of culture media, macrophage ontogeny, or host antimicrobial factors such as nitric oxide and itaconate. Access of Mtb to host-derived lipids required the ESX-1 secretion system and was inversely correlated with host triacylglycerol (TAG) synthesis. Inhibition of TAG formation in human macrophages partially restored Mtb ILI formation, revealing a metabolic gate that governs lipid flow between host lipid droplets and intracellular Mtb. Together, these findings establish a cross-species framework for decoding host-driven bacterial metabolic states and identify a key barrier limiting Mtb’s access to host lipid stores in human macrophages.

## INTRODUCTION

*Mycobacterium tuberculosis* (Mtb), the causative agent of tuberculosis (TB), remains one of the world’s leading infectious killers. Mtb’s ability to persist within macrophages is central to its success as a pathogen, yet the physiological states it adopts in different host environments remain incompletely defined. Mouse models have been indispensable for uncovering fundamental principles of Mtb–host interactions because of their genetic tractability, experimental reproducibility, and suitability for *in vivo* studies. However, it is well appreciated that key immunologic and metabolic features differ between murine and human macrophages, raising questions about how these differences shape Mtb physiology.

Comparative studies of species differences have largely focused on divergent host responses to infection rather than on the consequences for the bacterium itself. Two canonical examples, *nos2*, encoding inducible nitric oxide synthase, and *irg1*, which produces the antimicrobial metabolite itaconate, illustrate how the macrophage’s antimicrobial landscape differs across species (<u>Nair et al., 2018</u>; <u>Ohno et al., 2003</u>; <u>Shi et al., 2022</u>; <u>Voskuil et al., 2003</u>). Both are robustly induced in mouse macrophages but are more limited in human cells, resulting in distinct patterns of immune pressure. Yet how Mtb adapts metabolically and transcriptionally to these species-specific environments remains poorly understood.

Beyond direct antimicrobial stress, host cells impose nutritional constraints that strongly influence Mtb survival. Nutrient limitation represents a major axis of bacterial vulnerability, and Mtb’s capacity to remodel its metabolism under such conditions is a major determinant of persistence. Mtb’s strategies for nutrient acquisition, including fatty acids and cholesterol uptake, have therefore emerged as central to its pathogenesis and as potential therapeutic targets. For example, loss of the Mce4 transporter impairs cholesterol utilization and restricts growth in activated mouse macrophages (<u>Pandey & Sassetti, 2008</u>), underscoring how metabolic interplay between host and pathogen dictates infection outcomes.

Here, we apply a quantitative, cross-species framework to directly compare Mtb physiology in human and mouse macrophages. This analysis reveals extensive differential gene expression, highlighting distinct bacterial metabolic programs across hosts. Functional and metabolic validation identified a striking difference in the formation of intracellular lipid inclusions (ILIs), neutral lipid stores associated with metabolic quiescence and stress tolerance (<u>Garton et al., 2002</u>; <u>Mallick et al., 2021</u>) in Mtb. We further show that modulation of triacylglycerol biosynthesis in human macrophages alters Mtb ILI formation, defining a metabolic coupling that links host lipid flux to bacterial lipid storage.

These findings demonstrate that host species-specific environments drive distinct bacterial physiological states and uncover a previously unrecognized connection between host lipid droplets and Mtb lipid inclusions. More broadly, our cross-species approach provides a framework for dissecting how host context shapes pathogen state, offering a path toward more predictive models of Mtb pathogenesis.

## RESULTS

### Host species shape *M. tuberculosis* transcriptional programs in macrophages

Macrophages constitute the principal cellular niche for *Mycobacterium tuberculosis* (Mtb) and simultaneously mediate antimicrobial control. Foundational insights into host–Mtb interactions have come from *in vitro* infection models in which macrophages are analyzed for immunologic or microbiologic analyses. Mouse macrophages are widely used, yet several studies have underscored the limits of extrapolating antimicrobial mechanisms from mice to humans. For example, the contribution of reactive nitrogen species remains debated, and recent work points to species differences in the antimicrobial metabolite itaconate arising from a human-specific variant in aconitate decarboxylase 1. These observations highlight a gap in our understanding of how macrophage models behave across species in the context of Mtb infection.

Most cross-species comparisons have profiled the host response to defined stimuli (e.g., Toll-like receptor ligands) or infection, revealing differences in the magnitude and quality of host pathways (<u>Ahmed et al., 2020</u>; <u>Gilbertson et al., 2022</u>; <u>Schroder et al., 2012</u>). Here, we instead asked how Mtb changes state in distinct host macrophage environments. Measuring the Mtb transcriptome has proven informative for comparing macrophage activation states and Mtb genetic perturbations (<u>Pisu et al., 2020</u>; <u>Schnappinger et al., 2003</u>); we applied the same strategy across species.

We infected murine bone marrow-derived macrophages (BMDM) and human monocyte-derived macrophages (hMDM) and harvested RNA at 24 hours post-infection (*Materials and Methods*). Because bacterial transcripts are a minor fraction of total RNA in infected cells, we employed an optimized targeted-capture protocol to enrich for Mtb transcripts prior to sequencing (Figure 1A) (<u>Gunnarsson et al., 2025</u>). We observed that ∼20% of Mtb genes were differentially expressed (*p*_adj_ < 0.05) between mouse and human macrophages (Figure 1B), with approximately equal numbers higher in human versus mouse cells. Genes with higher expression in human cells included *papA3*, *mmpl8*, *pks4*, *esxP*, and *PPE51*; those higher in mouse cells included *icl1*, *hsp*, *lat*, and *whiB7* (Figure 1B and Supplemental Table 1).

**Figure 1.**
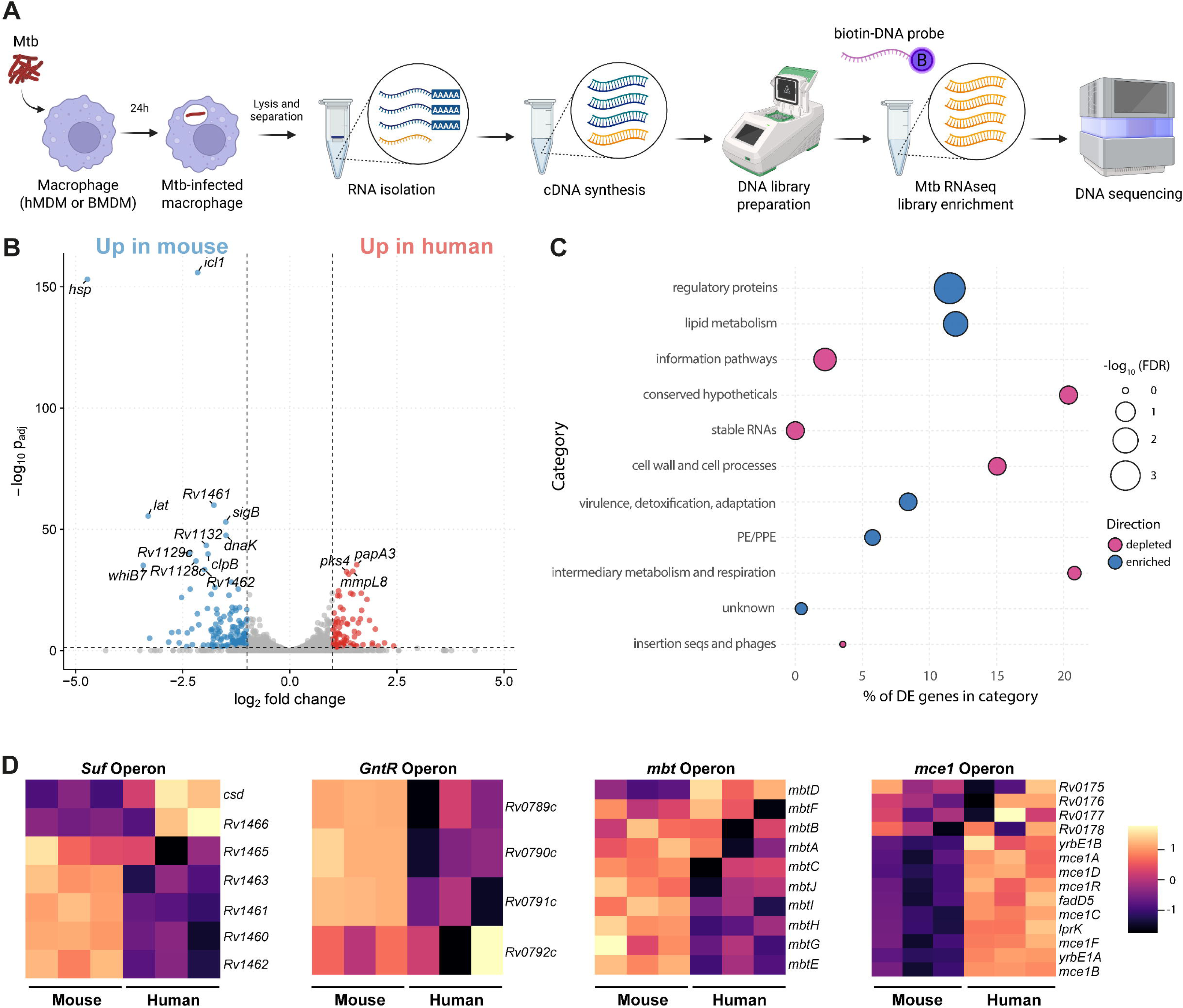
RNA seq analysis of differentially expressed genes in Mtb during human vs mouse macrophage infection. **(A)** Schematic of intracellular bacterial RNA sequencing workflow. Primary human monocyte-derived macrophages (hMDMs) and mouse bone marrow-derived macrophages (BMDMs) were infected with *M. tuberculosis* (Mtb) H37Rv wild-type (wt) during 24h. Cells were lysed, and bacteria and macrophages contents were separated by differential centrifugation and then combined for RNA isolation. Afterwards, cDNA was synthesized and amplified. Hybrid selection was done to enrich bacterial transcripts by using biotinylated-DNA probes complementary to the Mtb H37Rv transcriptome, excluding ribosomal and transfer RNA, before performing sequencing. **(B)** Volcano plot showing differentially expressed genes (DEGs) in Mtb during infection of human versus mouse macrophages. **(C)** Pathway enrichment analysis of Mtb DEGs grouped by functional category, depicting enriched and depleted genes during macrophage infection. **(D)** Heat maps showing Mtb operons differentially expressed during infection of mouse and human macrophages, including Fe–S complex assembly (*Suf* operon), reactive oxygen species response (*GntR* operon), mycobactin synthesis (*Mbt* operon) and fatty acid transport (*Mce1* operon).

Pathway-level analysis using *p*_adj_ < 0.05 and |log₂FC| > 1 indicated enrichment of genes involved in regulation and lipid metabolism among the differentially expressed set (Figure 1C). These trends were consistent when relaxing the fold change requirement. Enriched regulators included *prpR*, *mce1R*, and *tcrX*, and lipid-associated genes included *pks1–4*, *papA1–4*, and *umaA*. Motivated by these signals, we examined expression of a curated lipid metabolism gene set (fatty acid-responsive genes, transporters, and cholesterol-catabolism genes). We observed elevated expression of stress-response genes such as iron-stress (e.g., *Rv1460*–*66* and *csd*; which belong to the *Suf* operon), reactive oxygen species response (e.g., *Rv0789c*–*92c*; included in the *GntR* operon), as well as mycobactin biosynthesis genes (*mbtB*, *mbtD*, *mbtE*, *mbtI*; part of the *Mbt* operon) in mouse macrophages, whereas components associated with fatty acid import (e.g., *YrbE1A*, *YrbE1B*, *mce1A*, *mce1B*; which belong to the *Mce1* operon) were more highly expressed in human macrophages (Figure 1D). It is well known that lipids are very important for Mtb survival and replication during host infection and during axenic growth. Mtb can replicate inside lipid-loaded foamy macrophages (<u>Peyron et al., 2008</u>), and possesses the unique capacity to import and utilize host-derived fatty acids and cholesterol to synthesize its complex and lipid-rich cell envelope (<u>Gago et al., 2018</u>). In fact, host-derived lipids are the primary carbon source for Mtb *in vivo*, which are catabolized to fuel central metabolic pathways required for Mtb’s persistence (<u>Russell et al., 2009</u>). In the present work, we show that lipid import and metabolism are enriched in Mtb during macrophage infection, and that macrophages from different host species impose distinct microenvironmental pressures that drive Mtb into separable transcriptional states, with prominent differences in metabolic pathways and regulatory circuits.

### *M. tuberculosis* forms intracellular lipid inclusions in resting murine macrophages but not in resting human macrophages

We sought to orthogonally validate the transcriptomic differences between human and mouse macrophages with a focus on the lipid metabolism signal. As a high-throughput means to monitor lipid storage, we used a previously established lipid labeling protocol in which neutral lipids are labeled with BODIPY 493/503 (Figure 2A). BODIPY 493/503 is a lipophilic dye that partitions into neutral lipids such as triacylglycerol (TAG) and cholesteryl esters (CE). Earlier studies using this protocol initially utilized Mtb Erdman, so we first infected primary hMDM or BMDM with Erdman that constitutively expresses a cytosolic mCherry for 24 hours. Then, cells were fixed, stained with BODIPY 493/503, and imaged by confocal microscopy (Figure 2B). Strikingly, BODIPY-stained puncta were observed in the cytosol of intracellular Mtb during infection of mouse macrophages, but these were not present when infecting human macrophages (Figure 2B). Mtb can import neutral lipids from the host and accumulate them in cytosolic lipid droplets (LDs), best known as intracellular lipid inclusions (ILIs) (<u>Garton et al., 2002</u>). Similar to eukaryotic LDs, mycobacterial ILIs also contain a neutral lipid core comprised primarily of TAGs surrounded by a single phospholipid monolayer (<u>Mallick et al., 2021</u>). Accumulation of ILIs in Mtb leads to inhibition of bacterial replication and transition to a non-replicating persistence (NRP) state, and more importantly ILIs may contribute to the survival, reactivation and/or transmission of these bacteria following infection (<u>Garton et al., 2002</u>; <u>Mallick et al., 2021</u>). We next used an alternative protocol employing BODIPY palmitate (C16), a fluorescent fatty acid analogue that can be utilized to dynamically track lipid trafficking to Mtb (<u>Nazarova et al., 2018</u>; <u>Podinovskaia et al., 2013</u>). In this experiment, mCherry-expressing Mtb were used to infect macrophages. After specific infection periods, BODIPY C16 is added to culture media for a defined period of time (pulse) and followed by a one hour incubation without BODIPY C16 (chase). Using BODIPY C16 as a probe of dynamic lipid trafficking, we observed ILIs in BMDM but not in hMDM. (Figure 2B,C). These species differences extended to murine macrophage cell lines, including immortalized BMDMs (iBMMs) and RAW264.7 cells, which both displayed ILIs (Supplemental Figure 1). We therefore used iBMMs going forward to take advantage of an existing panel of macrophage mutants. As BODIPY C16 facilitated better ILI visualization than BODIPY 493/503, we opted to use BODIPY C16 for the remainder of our experiments. Taken together, these experiments provide visual validation that differential expression of lipid metabolic genes during murine and human macrophage infection leads to changes in lipid handling in Mtb.

**Figure 2.**
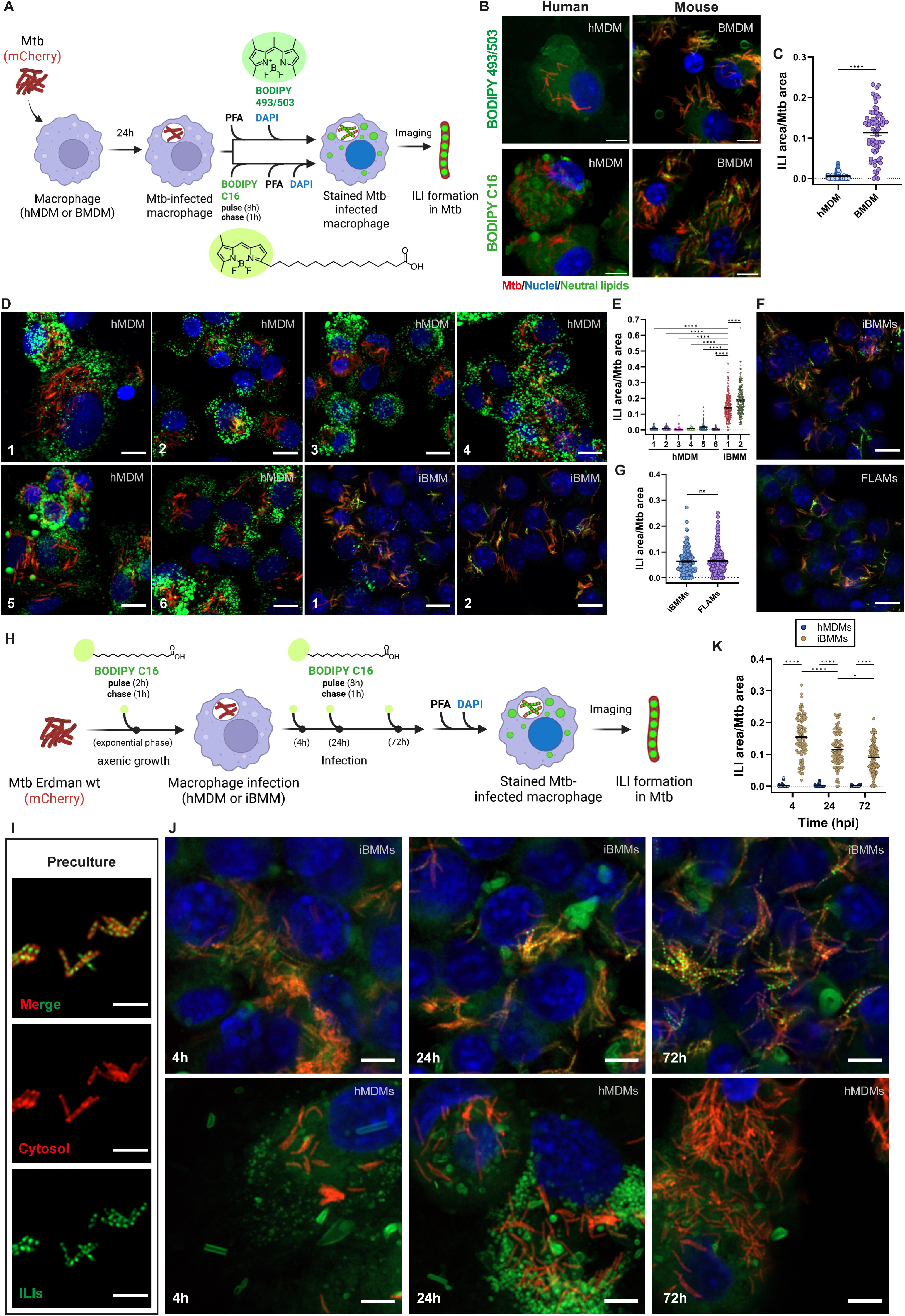
Intracellular lipid inclusions (ILIs) formed by Mtb during macrophage infection are species dependent. **(A)** Primary human macrophages (human monocyte-derived macrophages, hMDM) and primary mouse macrophages (bone marrow-derived macrophages, BMDM) were infected with cytosolic mCherry-expressing *M. tuberculosis* (Mtb) Erdman wild type (wt) for 24h. In one condition, samples were fixed with paraformaldehyde (PFA), stained with DAPI (nuclei), and neutral lipids labeled with BODIPY 493/503. In the other, neutral lipids were labelled by an 8 h pulse with fluorescent palmitate (BODIPY C16) followed by a 1h chase. Samples were imaged by Airyscan fluorescence microscopy **(B)**. Representative Airyscan images showing mCherry-labeled Mtb (red), nuclei (blue), and neutral lipids (green). Scale bars, 5 µm. **(C)** Quantification of BODIPY C16 fluorescence signal from Mtb ILIs in (B) using Cell Profiler; data are presented as ILI area / Mtb area ± SEM (N = 3). **(D)** hMDMs were cultured in the following media before infection: (1) RPMI + 10% FBS (R10) + M-CSF, (2) R10 + GM-CSF, (3) RPMI + 10% HS + M-CSF, (4) human plasma-like medium (HPLM) + 10% FBS + M-CSF, (5) HPLM + 10% HS + M-CSF, and (6) DMEM + 10% FBS + M-CSF. Immortalized BMDMs (iBMMs) were cultured in (1) DMEM + 10% FBS or (2) R10. Cells were infected for 24 h (MOI = 3 for hMDMs; MOI = 10 for iBMMs), subjected to an 8 h BODIPY C16 pulse and 1 h chase, fixed, stained with DAPI, and imaged by Airyscan fluorescence microscopy. Scale bars, 10 µm. **(E)** Quantification of ILI fluorescence from (D); data represent ILI area / Mtb area ± SEM (N = 3). **(F)** Murine immortalized macrophagess (iBMMs) and fetal liver-derived alveolar macrophages (FLAMs) were infected with cytosolic mCherry-expressing Mtb Erdman wt for 24 h, labeled with BODIPY C16 (8 h pulse, 1 h chase), fixed, and imaged as above. Scale bars, 10 µm. **(G)** Quantification of ILI fluorescence from (F); data represent ILI area / Mtb area ± SEM (N = 3). **(H)** Cytosolic mCherry-expressing Mtb Erdman wt was cultured in 7H9^OADC^ and, at exponential phase, labeled with BODIPY C16 (2 h pulse, 1 h chase). An unlabeled aliquot of the same preculture was used to infect iBMMs or hMDMs. Infections were analyzed at 4, 24, and 72 h post-infection using an 8 h BODIPY C16 pulse and 1 h chase. **(I)** Airyscan image of Mtb preculture showing cytosolic mCherry (red) and neutral lipids (green). Scale bars, 2 µm. **(J)** Airyscan images of Mtb-infected iBMMs (top) or hMDMs (bottom) showing mCherry (red), nuclei (blue), and neutral lipids (green). Scale bars, 5 µm. **(K)** Quantification of ILI fluorescence from (J); data represent ILI area / Mtb area ± SEM (N = 3). For hMDM data, individual colors denote independent donors. All data represent a minimum of three independent experiments.

We next performed additional experiments to confirm that the differences in ILI formation observed were genuine and not artifacts of media composition. Murine macrophages are commonly cultured in a media base of Dulbecco’s Modified Eagle Medium (DMEM) while human macrophages are commonly cultured in Roswell Park Memorial Institute (RPMI) 1640 media. Alternative culture media such as human plasma-like media (HPLM) can be used. Human macrophage culture media are often supplemented with growth factors such as M-CSF or GM-CSF to support differentiation. Furthermore, as an alternative to the use of heat-inactivated fetal bovine serum (FBS) in culture media, heat-inactivated human serum (HS) can be used. To test the hypothesis that the differences in ILI detection in human versus mouse macrophages are attributable to species differences and not protocol differences, human macrophages were cultured in the following media: (1) RPMI + 10% FBS (R10) + M-CSF, (2) R10 + GM-CSF, (3) RPMI + 10% HS + M-CSF, (4) HPLM + 10% FBS + M-CSF, (5) HPLM + 10% HS + M-CSF, (6) DMEM + 10% FBS + M-CSF prior to ILI analysis. Conversely, iBMMs were cultured in either (1) DMEM + 10% FBS or (2) R10. We next repeated our BODIPY C16 pulse-chase experiments and were able to detect ILIs in iBMMs under all culture conditions. ILIs remained undetectable in hMDMs in all culture conditions (Figure 2D,E). These results further confirmed the differences in Mtb metabolic state between human and murine macrophages.

To determine whether the ILI phenomenon observed in murine macrophages extended to macrophages from different ontogenies, we evaluated murine fetal liver-derived macrophages (FLAMs) as a distinct lineage model. We compared iBMMs and FLAMs following infection with Mtb and performed the BODIPY C16 pulse-chase experiment. We readily detected ILIs in both iBMMs and FLAMs (Figure 2F,G).

It is possible that the observed species difference is due to distinct kinetics in ILI formation. To test this, we determined how soon after Mtb infection this differential ILI phenotype is observed. We grew Mtb Erdman expressing cytosolic mCherry in 7H9 medium supplemented with glycerol, Tween and OADC (oleic acid, bovine albumin, dextrose, and catalase). At exponential phase, the capacity for ILI formation was tested by a 2h BODIPY C16 pulse-1h chase. This same unlabeled Mtb culture was used to infect either hMDMs or iBMMs. After 4, 24 or 72h of infection, BODIPY C16 pulse-chase experiments were performed (Figure 2H). ILIs were observed in the Mtb preculture during axenic growth (Figure 2I). This ILI competent trait in Mtb was conserved throughout axenic growth in complex media (Supplemental Figure 2). Surprisingly, immediately upon hMDM infection, Mtb was devoid of ILIs and the capacity for ILI formation was not restored at the later infection timepoints. In contrast, during iBMM infection, despite being dimmer right after infection, ILIs were readily formed throughout the course of the infection (Figure 2J,K). Taken together, these results indicate that access to lipids by intracellular Mtb for ILI formation is different in these two species.

### Knockout of host factors with well-characterized species differences does not affect ILI formation in *M. tuberculosis* during macrophage infection

Previous studies have identified many host factors that are differentially regulated in human versus mouse macrophages. We therefore tested if these factors accounted for the observed ILI formation in murine versus human macrophages (<u>Gilbertson et al., 2022</u>; <u>Mills et al., 2018</u>; <u>Schroder et al., 2012</u>; <u>Zhang et al., 2013</u>). We focused on nitric oxide and itaconate as candidate molecules. The field continues to debate the role of nitric oxide synthase 2 (NOS2) in antimicrobial programs in human cells though recent studies suggest that the genomic architecture of the human NOS2 locus contributes to distinct expression patterns between humans and mice (<u>Cooper et al., 2000</u>; <u>Gilbertson et al., 2022</u>). The enzyme responsible for itaconate production, *cis*-aconitate decarboxylase 1 (ACOD1), encoded by the gene *irg1*, has a point mutation in humans that reduces itaconate production by orders of magnitude compared to the murine enzyme (<u>Chen et al., 2019</u>; <u>Michelucci et al., 2013</u>; <u>Shi et al., 2022</u>). Both nitrosative species and itaconate have been implicated in regulation of Mtb metabolism (<u>Cooper et al., 2000</u>; <u>Nair et al., 2018</u>; <u>Ohno et al., 2003</u>; <u>Priya et al., 2025</u>; <u>Voskuil et al., 2003</u>). We used CRISPR-Cas9 to knock out *irg1* or *nos2* in murine macrophages. After confirming knockout by Western blotting, we again performed our BODIPY C16 pulse-chase experiments in these macrophages infected with Mtb (Figure 3). Neither deletion of *irg1* (Figure 3A,B) nor *nos2* (Figure 3C,D) was able to disrupt ILI formation.

**Figure 3.**
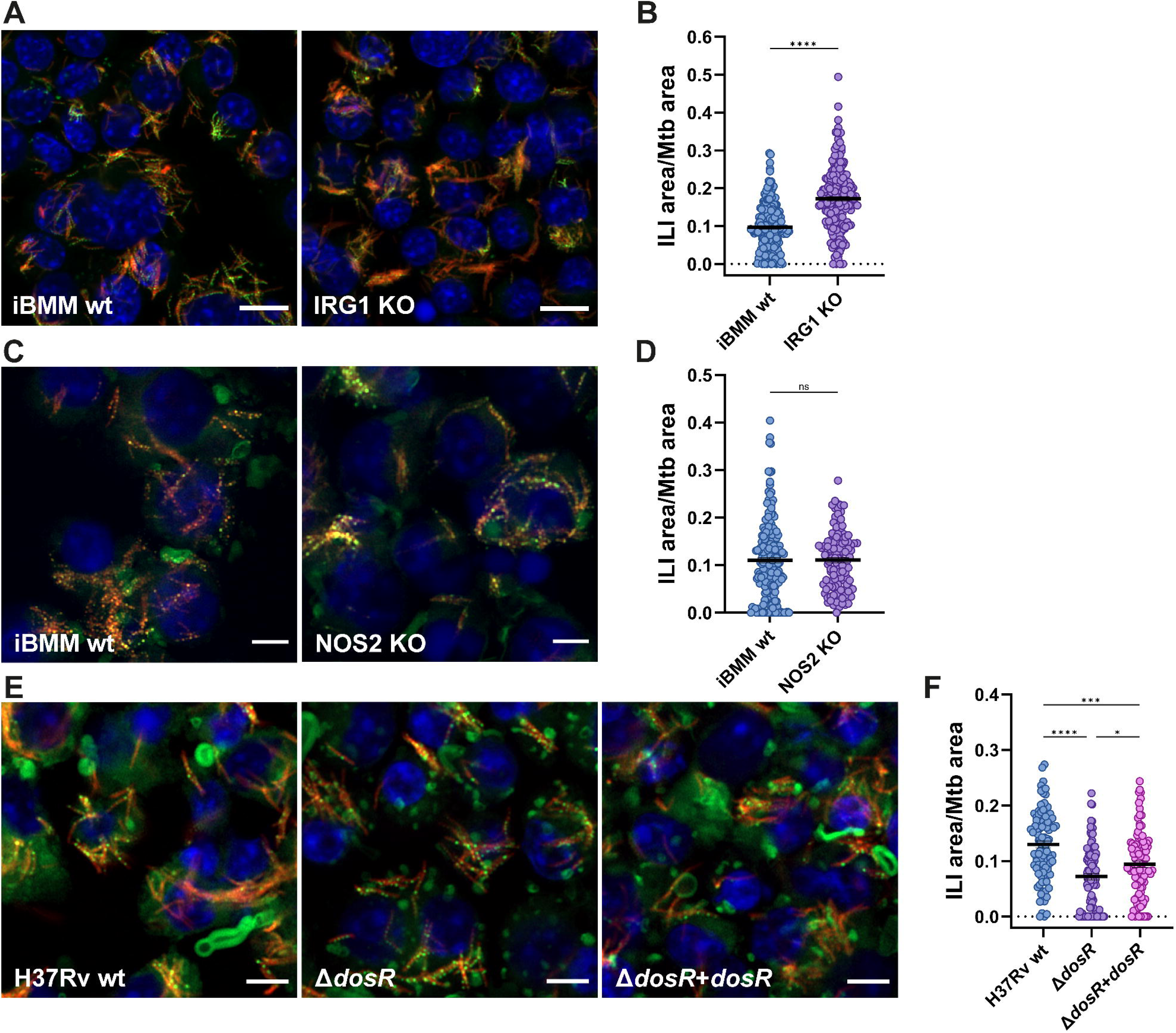
Knockout of mammalian and bacterial factors have different effects on ILI formation in *M. tuberculosis* in macrophages. **(A–D)** Immortalized murine bone marrow–derived macrophages (iBMMs) expressing Cas9 or knockout (KO) mutants for *irg1* (A) and *nos2* (C) were infected for 24 h with cytosolic mCherry-expressing Mtb Erdman. Following infection, neutral lipids were labelled with BODIPY C16 (8 h pulse, 1 h chase), cells were fixed, nuclei were stained with DAPI, and samples were imaged using Airyscan fluorescence microscopy (A,C). (B, D) Quantification of ILI signal from (A and C) using CellProfiler. Data are shown as ILI area / Mtb area ± SEM (N = 3). **(E, F)** iBMMs were infected with Mtb H37Rv wild type (wt), Δ*dosR*, or complemented Δ*dosR* + *dosR* strains for 24 h, BODIPY C16 pulse-chase was performed and samples were treated and imaged as above (E). (F) Quantification of ILI signal from (E) using CellProfiler. Data are shown as ILI area / Mtb area ± SEM (N = 3). Infections were conducted at MOI = 10. Scale bars, 5 µm.

### Deletion of bacterial genes involved in dormancy or virulence have different effects on ILI formation

Previous studies have argued that ILI formation in axenic culture occurs in response to hypoxia or other stresses associated with dormancy. The transcriptional regulator DosR has been implicated in regulation of these stress responses (<u>Mehra et al., 2015</u>; <u>Park et al., 2003</u>). We obtained an Mtb mutant lacking *dosR* as well as its complemented strain, and generated fluorescent versions of both (<u>Park et al., 2003</u>). We next infected iBMMs with these strains and performed our BODIPY C16 pulse-chase experiments. Deletion of *dosR* did not disrupt ILI formation in Mtb during murine macrophage infection, consistent with previous predictions by others (<u>Fines et al., 2023</u>) (Figure 3E,F). In our experiments, BODIPY C16 was added extracellularly and thus must be taken up by the macrophage and then trafficked to the intracellular bacterium. We hypothesized that Mtb mutants that influence phagosome integrity might impact BODIPY C16 acquisition during infection. We generated a fluorescent EccCa1 transposon mutant (*eccCa1:Tn*) and next performed our BODIPY C16 pulse-chase experiments. EccCa1 is an ATPase that regulates activity of the ESX-1 secretion system (<u>Luthra et al., 2008</u>). The ESX-1 system exports proteins that contribute to phagosomal membrane damage (<u>Groschel et al., 2016</u>). Strikingly, during iBMM infection, the *eccCa1:Tn* mutant lacked detectable ILIs (Supplemental Figure 3A,B). We next asked whether loss of ESX-1 activity disrupted BODIPY C16 acquisition during axenic growth, to assess whether the phenotype was specific to macrophage infection or reflected a general defect in lipid import. In contrast to the loss of ILI signal in iBMMs observed with the ESX-1 mutant, we observed no defect in lipid import by the *eccCa1:Tn* mutant in axenic culture (Supplemental Figure 3C). This strongly suggests that the ESX-1 system is required for ILI formation by the bacteria during macrophage infection, in addition to its canonical roles in phagosomal membrane disruption and virulence effector secretion.

### Impaired triacylglycerol synthesis in human macrophages permits ILI formation in infecting *M. tuberculosis*

While our experiments with Mtb mutants expanded our understanding of bacterial factors influencing lipid acquisition in murine macrophages, the factors contributing to the observed interspecies differences in Mtb ILI formation remained unclear. Our data did not indicate a defect in BODIPY C16 import by hMDMs as host lipid droplets (LDs) were readily loaded with BODIPY C16 (Figure 2). The rate-limiting enzyme involved in TAG synthesis in mammalian cells is diacylglycerol acyltransferase 1 (DGAT1), which catalyzes the synthesis of TAG from diacylglycerol (DAG) and long-chain fatty acyl-CoAs (<u>Yen et al., 2008</u>). To examine the role of TAG synthesis and host LD formation in Mtb ILI formation, human macrophages were pre-treated with two DGAT1 inhibitors (T863 and Pradigastat) following a previously established protocol linking LD formation to the efficacy of the lipophilic antibiotic bedaquiline (<u>Greenwood et al., 2019</u>). hMDMs were pre-treated for 48 hours with vehicle or drug prior to infection with mCherry Mtb (Figure 4A). Cells were infected with Mtb in the absence of extracellular drug for 4 hours. After washing away extracellular Mtb, medium containing vehicle or drug was added back for the remainder of the experiment. 24 hours following infection, we performed our BODIPY C16 pulse-chase experiment and visualized samples by fluorescence microscopy (Figure 4B). We first confirmed that drug treatment decreased the formation of host LDs by confocal microscopy (Figure 4C,D). We then quantified ILI formation as a function of drug and concentration and found that inhibition of host TAG synthesis resulted in detectable ILI signal in Mtb (Figure 4E). The enhanced Mtb ILI signal in hMDMs was dose-dependent and occurs with both drugs. Notably, this inhibition did not result in detectable ILI signal in all intracellular Mtb, suggesting that host TAG synthesis may be one component of regulation of Mtb metabolic state. We next tested whether modulating Mtb ILI formation in hMDMs could affect bacterial burden. Using Mtb area as a proxy for bacterial burden, we tested whether restoring ILIs in hMDMs through inhibition of TAG synthesis altered bacterial load; however, we observed no significant change (Figure 4F). These findings suggest that fatty acids stored as TAGs in host LDs are not readily available to Mtb for ILI formation in hMDMs. Instead, host LDs may sequester lipids away from the bacteria, or imported lipids may be preferentially used for cell wall synthesis required for bacterial elongation and replication.

**Figure 4.**
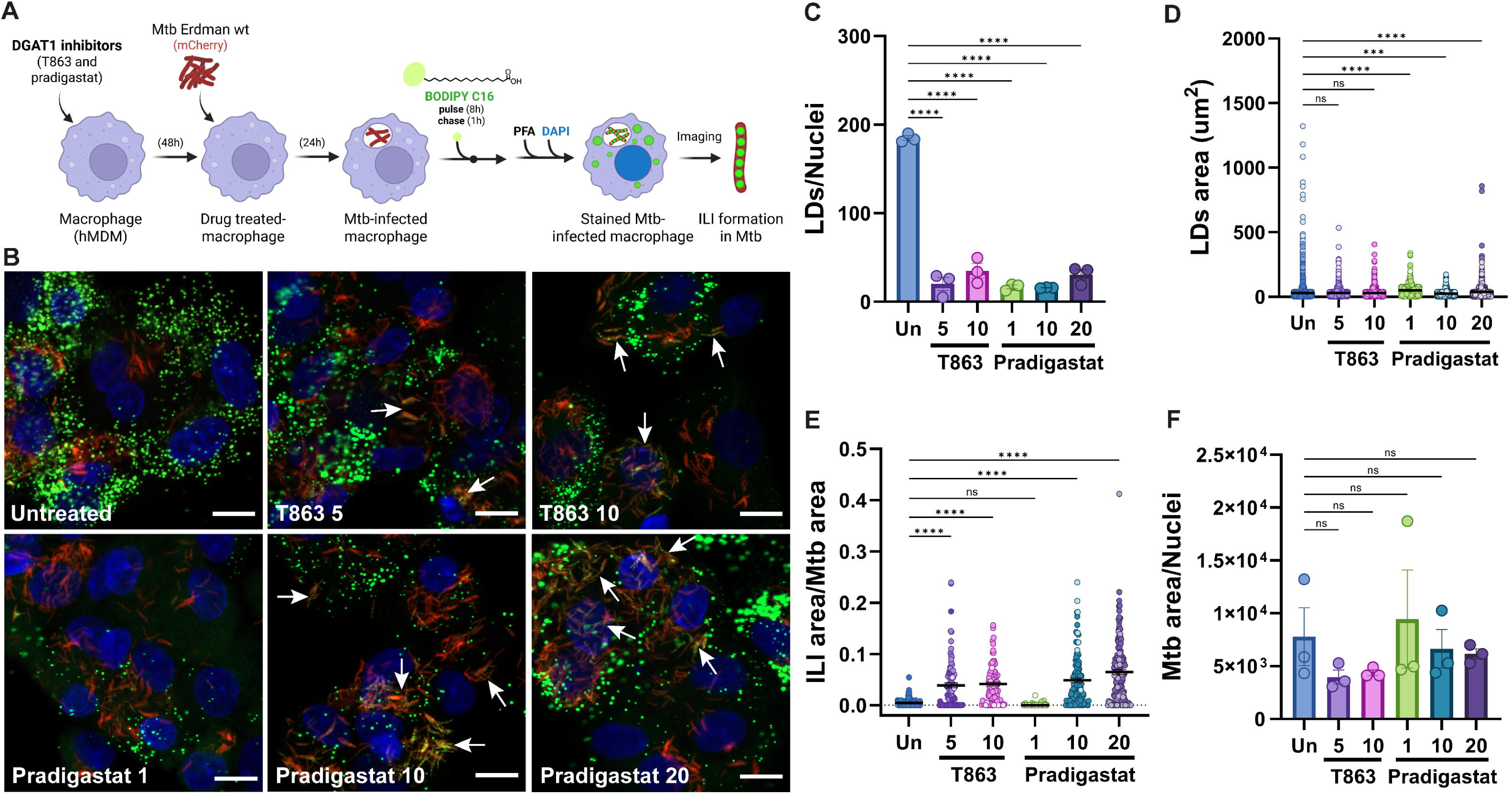
Inhibiting host TAG synthesis with DGAT1 inhibitors permits intracellular lipid inclusion (ILI) formation in Mtb in human macrophages. **(A, B)** Human macrophages (hMDMs) were pre-treated for 48h with two distinct diacylglycerol acyltransferase 1 (DGAT1) inhibitors—T863 (5 or 10 mg/L) or Pradigastat (1, 10, or 20 mg/L)—and then infected for 24h with cytosolic mCherry-expressing Mtb Erdman at MOI = 3. Following infection, neutral lipids were labelled with BODIPY C16 (8 h pulse, 1 h chase), cells were fixed, nuclei were stained with DAPI, and samples were imaged using Airyscan fluorescence microscopy. White arrows indicate Mtb containing ILIs. Scale bars, 10 µm. **(C–E)** Quantification of (C) lipid droplet (LD) number per nucleus, (D) LD area (µm²), and (E) ILI area per Mtb area from images in (B). Data are presented as mean ± SEM (N = 3) (C) or as ILI area / Mtb area ± SEM (N = 3) (D,E). **(F)** Bacterial burden represented as Mtb area per nucleus ± SEM (N = 3). Points are color-coded to denote individual human donors.

## DISCUSSION

As an intracellular pathogen, *Mycobacterium tuberculosis* (Mtb) resides primarily within macrophages, where it must adapt to host-imposed metabolic and immune constraints. Our cross-species analyses reveal that these pressures differ markedly between mouse and human macrophages, driving Mtb into distinct physiological states. In mouse macrophages, Mtb upregulates iron acquisition, oxidative stress, and lipid metabolism pathways including the *suf* and *mbt* operons whereas in human macrophages, the bacterium preferentially induces fatty acid import and β-oxidation machinery such as *mce1*, *fadD*, and *yrbE* genes (Figure 1). These transcriptional differences manifest in a striking phenotypic divergence: Mtb forms intracellular lipid inclusions (ILIs) during infection of mouse macrophages but not in human macrophages (Figure 2 and Supplemental Figure 1).

ILIs have been observed previously in sputum isolates and in macrophages under specific stress conditions (e.g., (<u>Knight et al., 2018</u>)), but their biological origin and regulation have remained incompletely understood. Our data reveal that ILI formation is not an invariant feature of Mtb infection but rather a host species-specific outcome reflecting distinct intracellular environments. Multiple stressors such as hypoxia, nutrient limitation, or redox stress can promote triacylglycerol (TAG) accumulation in Mtb (<u>Daniel et al., 2011</u>; <u>Garton et al., 2002</u>; <u>Lee et al., 2013</u>; <u>Peyron et al., 2008</u>), yet our findings demonstrate that even in the absence of such stimuli, infection of mouse macrophages alone is sufficient to trigger ILI formation. In contrast, human macrophages do not elicit this response, underscoring that Mtb’s metabolic state is tightly coupled to host-specific cues (Figure 2 and Supplemental Figure 1). ILI formation is also observed in Mtb during axenic growth in complex media (Supplemental Figure 2).

Mtb possesses respiratory flexibility, which allows it to grow optimally in aerobic conditions but also survive under hypoxia, still maintaining bioenergetic homeostasis. Mtb is considered a prototrophic bacterium, which possesses a broad metabolic repertoire and a unique lipid-rich and complex cell envelope that provide the pathogen the ability to survive within diverse microenvironments during host cell infection (<u>Warner et al., 2025</u>). The Mtb cell envelope represents a highly hydrophobic and multilayered dynamic barrier, whose biophysical structural properties may also impact host infection (<u>Batt et al., 2024</u>). The ancient co-evolution of Mtb with its hosts has imposed selective pressures that have shaped, and continue to influence, its physiology and metabolism (<u>Comas et al., 2013</u>; <u>Wirth et al., 2008</u>). Mtb’s ability to co-catabolize both carbon and nitrogen sources, as well as to modify its responses to different host cell types magnifies its metabolic flexibility and augments its success as a pathogen (<u>Borah et al., 2021</u>). Taken together, Mtb’s transcriptional plasticity allows it to respond to diverse host environmental cues, and it denotes that the bacterial metabolic state is a major driver of its physiology and resistance to the host antimicrobial responses.

In the present study, we found that ILI formation in Mtb does not depend on host nitric oxide or itaconate production (Figure 3), two antimicrobial metabolites that differ between mouse and human macrophages, suggesting that other, as yet undefined host signals govern this process. Instead, our data implicate host neutral lipid metabolism as a key determinant. Inhibition of host TAG synthesis through DGAT1 blockade unexpectedly enabled Mtb to access host lipids and restored ILI formation (Figure 4), suggesting that sequestration of fatty acids within host lipid droplets (LDs) can modulate bacterial lipid metabolism. This observation highlights an underappreciated connection between host LD homeostasis and bacterial nutrient availability. It also suggests that the interplay between host TAG metabolism and bacterial lipid storage could influence the intracellular efficacy of lipophilic antimicrobials, as previously proposed (<u>Greenwood et al., 2019</u>).

We also identified bacterial determinants of ILI formation. While the DosR regulon—long associated with the hypoxia response (<u>Mehra et al., 2015</u>; <u>Park et al., 2003</u>)—was dispensable (Figure 3), the ESX-1 secretion system (<u>Groschel et al., 2016</u>) proved essential. Mtb mutants lacking ESX-1 failed to form ILIs and exhibited restricted lipid acquisition and replication within macrophages (Supplemental Figure 3). These findings extend the role of ESX-1 beyond virulence and phagosomal rupture to include access to host-derived lipids. We speculate that species-specific differences in ESX-1 activity or the degree of phagosomal permeabilization may contribute to the divergent lipid phenotypes observed in human and mouse macrophages.

During infection, Mtb relies primarily on host-derived lipids as carbon sources for energy production and to synthesize its own cell envelope. Formation of ILIs as a TAG storage bacterial organelle was first observed by Waltermann (<u>Waltermann et al., 2005</u>; <u>Waltermann & Steinbuchel, 2005</u>). Although TAG deposition within LDs is conserved in eukaryotic cells from plants, animals and yeast, ILI formation has only been observed in a small group of prokaryotes, including actinomycetes such as *Mycobacterium*, *Rhodococcus*, *Nocardia* and *Streptomyces* (<u>Waters & Eijkelkamp, 2024</u>). Host fatty acid uptake in Gram-positive pathogenic bacteria such as *Streptococcus pneumoniae* and *Staphylococcus aureus* is essential for membrane phospholipid synthesis. Also, in Gram-negative human pathogens such as *Salmonella typhimurium*, *Pseudomonas aeruginosa*, *Acinetobacter baumannii* and *Escherichia coli*, the utilization of host-derived fatty acids is key for bacterial virulence (<u>Waters & Eijkelkamp, 2024</u>). Given that *de novo* fatty acid synthesis by fatty acid synthase II (FAS II) is highly energetically demanding, some pathogenic bacteria have evolved to use exogenously acquired fatty acids only for membrane biogenesis and replication. Apart from *M. tuberculosis* and *A. baumannii*, it would be interesting to identify other pathogens that similarly depend on host fatty acids during infection, and to determine how host-derived cues modulate their metabolic states and promote storage of TAGs or other lipids within ILIs.

Collectively, our findings uncover a new axis of host–pathogen metabolic interaction in which ILI formation serves as a readout of Mtb’s access to host lipids. This process depends jointly on host TAG synthesis and bacterial ESX-1 function and varies fundamentally between species. While ILIs have traditionally been viewed as markers of dormancy, our results suggest they can also arise as a context-dependent adaptation to the metabolic landscape of the host cell. Future work should elucidate the molecular signals that trigger ILI formation in mouse macrophages, define whether similar conditions occur *in vivo*, and determine how modulation of host lipid metabolism or ESX-1 activity reshapes bacterial physiology. Understanding these species-specific interfaces between host and pathogen metabolism will be critical for identifying new therapeutic strategies to restrict Mtb survival in human macrophages.

## MATERIALS AND METHODS

### Bacteria and media

*Mycobacterium tuberculosis* (Mtb) H37Rv and Erdman strains were grown at 37°C in 7H9^OADC^ medium (Middlebrook 7H9 Broth supplemented with 0.2% glycerol, 0.05% Tween-80, and OADC (oleic acid, bovine albumin, dextrose, and catalase)) with shaking at 65 rpm. Antibiotics were added to the media when required, at the following concentrations: 20 µg/mL Zeocin (Thermo), for pBB84 plasmid-containing strains expressing a cytosolic mCherry.

### BODIPY stock preparation

For neutral lipid labelling during either bacterial axenic growth or during macrophage infection, a 5 mM stock of BODIPY palmitate (C16) (BODIPY FL C_16_, Thermo) was prepared in dimethyl sulfoxide (DMSO). The 5 mM BODIPY C16 stock was diluted in 1% fatty acid-free BSA prepared in PBS (Phosphate-Buffered Saline buffer solution), to obtain a 100 µM BODIPY C16 stock, as previously described (<u>Nazarova et al., 2018</u>). The 100 µM BODIPY C16 stock was vortexed until the solution turned green and kept at 37 °C wrapped in aluminum foil to protect it from light. For lipid incorporation analysis during axenic culture, at the time of labelling, the 100 µM BODIPY C16 stock was diluted in 7H9^OADC^ and vortexed to obtain an 8 µM working solution. For lipid incorporation analysis during macrophage infection, the 100 µM BODIPY C16 stock was diluted in the desired media to obtain an 8 µM BODIPY C16 working solution in media. Alternatively, a 5 mM stock of BODIPY 493/503 (Thermo) was prepared in DMSO and further diluted in PBS to obtain a working solution at 5 µM, which was used for neutral lipid staining post-fixation.

### BODIPY C16 lipid labeling during axenic growth

Mtb strains were grown in 7H9^OADC^ until logarithmic phase (OD ≤1) and a 0.5 mL aliquot was transferred to a 1.5 mL microcentrifuge tube. Samples were centrifuged at 3900 rpm for 5 min at room temperature. Then, the supernatant was removed, and the pellet was resuspended in 0.5 mL of 8 µM BODIPY C16 working solution in 7H9^OADC^. Samples were protected from light by wrapping the tube in foil and they were incubated at 37 °C for 2h with shaking (pulse). After the pulse, samples were centrifuged at 3900 rpm for 5 min at room temperature. The supernatant was removed, and the pellet was resuspended in 0.5 mL of 7H9^OADC^ and incubated again at 37 °C for 1h with shaking (chase). After the chase, samples were centrifuged at 3900 rpm for 5 min at room temperature, supernatant was removed, and samples were fixed by resuspending the pellet in 0.5 mL of 4% paraformaldehyde (PFA) in PBS. After fixation, two series of washing steps with 0.5 mL of PBS were done, and samples were resuspended milli-Q water, plated and visualized by high-resolution fluorescence microscopy.

### Human macrophage differentiation and culture

For human cell isolation and differentiation, de-identified, truly anonymous buffy coats were obtained from the Massachusetts General Brigham Hospital. Peripheral blood mononuclear cells (PBMCs) were isolated by density-based centrifugation with Ficoll (GE Healthcare). CD14+ monocytes were further isolated using a CD14-positive selection kit (Stemcell). Isolated monocytes were differentiated to macrophages using a variety of media conditions.

1. Buffy media (Phenol-free RPMI 1640 (Gibco) supplemented with 10% heat-inactivated fetal-bovine serum (HI-FBS) (Gibco), 10 mM HEPES (Corning), and 2 mM L-glutamine (Sigma) along with 25 ng/mL of either M-CSF or GM-CSF (Biolegend), depending on the experimental design).
2. HPLM media (Human Plasma Like Media (Invitrogen) supplemented with 10% HI-FBS (Gibco), 10 mM HEPES (Corning), and 2 mM L-glutamine (Sigma) along with 25 ng/mL of M-CSF (Biolegend).
3. BMDM media (DMEM (Gibco) supplemented with 10% HI-FBS (Gibco), 10 mM HEPES (Corning), and 2 mM L-glutamine (Sigma) along with 25 ng/mL of M-CSF (Biolegend).

In some cases, heat-inactivated human AB serum was used instead.

Cell differentiation was carried out by plating the cells on ultra low-attachment plates or in 12-well chamber slides (IBIDI). In the case of cells plated in ultra low-attachment plates, after 6 days of differentiation, cells were detached using PBS supplemented with 2mM EDTA and were replated in 12-well chamber slides at a cell density of 2×10^5^ cells/well. The cells were allowed to re-adhere overnight prior to infection.

### Bone marrow-derived macrophage (BMDM) isolation and differentiation

Femurs and tibias were harvested from C57BL/6 mice, stripped of muscle, and washed in ethanol and sterile PBS. Under sterile conditions, the ends of the bones were cut, and marrow was flushed into DMEM (Gibco) containing 10% HI-FBS (Bio-Techne) and 1% penicillin-streptomycin (Gibco) using a 27G needle and syringe. A single-cell suspension was achieved by repeatedly passing the marrow through the needle and filtering it through a pre-wetted 70 µm filter. Cells were pelleted and resuspended in 2 mL ACK (ammonium-chloride-potassium) lysis buffer (Gibco) for 1 minute on ice to lyse red blood cells. Following red blood cell lysis, cells were resuspended at 1.5×10^6^cells/mL in DMEM (Gibco) containing 10% HI-FBS (Bio-Techne), 1% penicillin-streptomycin (Gibco), and 25 ng/mL murine M-CSF (PeproTech). Two milliliters (3×10^6^ cells) were plated per well on a 6-well, tissue culture-treated dish. A 50% media replacement was performed at days 2 and 4 post-plating. Prior to Mtb infection, macrophages were washed with antibiotic-free media and all infections and post-infection experiments were performed in the absence of antibiotics.

### Murine macrophage cell line culture (iBMM, RAW, FLAMs)

For other murine macrophages such as immortalized BMDMs (iBMMs) and RAW264.7 macrophages (RAWs), D10 (DMEM supplemented with 10% HI-FBS, and 2mM L-glutamine) media was used. For culture of fetal liver-derived alveolar murine macrophages (FLAMs), (<u>Thomas et al., 2022</u>) supplemented with 10% HI-FBS, 30 ng/mL recombinant mouse GM-CSF and 20 ng/mL recombinant human TGFβ1 was used. In all cases, murine macrophages were plated at a cell density of 1.75×10^5^ cells/well in 12-well chamber slides.

### Preparing *M. tuberculosis* cultures for infection

Mtb strains were grown in 7H9^OADC^ until logarithmic phase (OD ≤1) and for each experiment a 3-5 mL aliquot was transferred to a 15 mL conical tube. Mtb culture was pelleted at 3900 rpm for 5 min at room temperature. Supernatant was removed and pellet was washed by resuspending it in an equal volume of PBS and centrifuging at 3900 rpm for 5 min at room temperature. Then, supernatant was removed and large Mtb clumps were allowed to settle down by centrifuging at 500 rpm for 5 min at room temperature. Afterwards, the supernatant was collected, and the OD_600_ was measured, using PBS as blank. Assuming an OD_600_ of 1 = 3e^8^ Mtb/mL, the volume needed for the desired MOI was calculated. hMDMs and mouse macrophages were infected to achieve approximately a 50% infection rate.

### BODIPY C16 lipid labeling during infection

For hMDM infection, an MOI of 3 was used, and for murine macrophages, an MOI of 10 was used. Mtb phagocytosis proceeded for 4h at 37 °C with 5% CO_2_. After phagocytosis, Mtb-containing media was removed from wells and two serial washing steps were performed using prewarmed PBS. Then, infections were incubated at 37 °C with 5% CO_2_ for 24-72h, depending on the experimental design.

To perform the BODIPY pulse-chase analysis: the 100 µM BODIPY C16 stock was diluted with the appropriate media to obtain an 8 µM BODIPY C16 working solution, and it was added to the appropriate wells. Samples were incubated for 8h at 37 °C with 5% CO_2_ (pulse), and then 8 µM BODIPY C16-containing media was removed from the wells and it was replaced with media only. Samples were incubated for 1h at 37 °C with 5% CO_2_ (chase). Afterwards, media was removed from the wells and samples were fixed with 4% PFA in PBS. Then, samples were washed twice with PBS, stained with DAPI, and chamber slides were mounted and visualized by high-resolution fluorescence microscopy.

### High-resolution fluorescence microscopy

For imaging Mtb grown axenically, PFA-fixed samples were washed twice with PBS and resuspended in Milli-Q water. Depending on the bacterial density of the samples, appropriate dilutions were prepared in Milli-Q water and samples were applied to 35 mm dishes with 14 mm diameter glass coverslips uncoated (Mattek). After the samples dried at room temperature, another set of dishes was prepared with 1% agarose in Milli-Q to mount the samples.

For imaging the Mtb-infected macrophages in 12-well chamber slides, PFA-fixed samples were washed twice with PBS, and half of each sample was treated with 60% isopropanol to permeabilize the cells. After DAPI staining, samples were washed twice with PBS and mounted using ProLong Diamond Antifade Mountant (Thermo) and VWR Coverslips 24×60mm No.1.5 (Avantor). Samples were allowed to cure for at least 24h before imaging. Both plates and chamber slides were imaged using confocal fluorescence microscopy using the Cell Discoverer 7 microscope (Zeiss) and the modular image acquisition software ZEN (Zeiss). The Alexa Fluor 568 (AF568) channel was used to acquire the mCherry signal, and the Alexa Fluor 488 (AF488) channel was used to acquire the green fluorescence from neutral lipids stained with BODIPY C16 or BODIPY 493/503. Images were acquired using the 20× and 40× objective lenses at a resolution of 2048 x 2048 pixels. Images were acquired with a 1.0 Airy unit pinhole. Images acquired by confocal or Airyscan microscopy were exported as .czi files, and analyzed using ImageJ and Cell Profiler software.

### RNA extraction and purification

Bacterial RNA was harvested from infected macrophages as previously described (<u>Pisu et al., 2020</u>). Infected macrophages were lysed in TRIzol (Thermo Fisher Scientific) and centrifuged at 10,000xg for 20 min at 4 °C to pellet intact bacteria. The supernatant, representing the host RNA fraction, was collected and filtered twice through a 0.2 µm PVDF filter to ensure pathogen inactivation. The bacterial pellet was resuspended in fresh TRIzol containing Lysing Matrix B beads (MP Biomedicals) and disrupted using a bead beater homogenizer (MP Biomedical) at 6.5 m/s for 45s, followed by a 5 min incubation on ice and a second 30 s bead-beating cycle. Following lysis, the host and bacterial RNA fractions were combined, then extracted with chloroform (200 µL per 1 mL TRIzol), and purified using on-column cleanup with DNase treatment (RNA Clean & Concentrate, Zymo Research).

### Library preparation and sequencing

200ng of total RNA was used as input for strand-specific library preparation with the Agilent SureSelect XT HS2 RNA Kit, following the manufacturer’s instructions, with a 1:2 adapter dilution, and an additional 0.8X/1.2X double SPRI post-ligation to deplete ribosomal RNA. For intracellular bacterial RNA sequencing, 200ng unpooled libraries were hybrid-selected for bacterial transcripts using a custom set of biotinylated probes complementary to the Mtb H37Rv transcriptome, excluding ribosomal and transfer RNAs, and probes were used at a 1:5 dilution to account for bacterial transcript being a low proportion of input libraries. All libraries were sequenced on an Illumina NextSeq 550 using 150 bp reads.

### Bioinformatic processing

Sequencing reads were processed using a previously published in-house pipeline for intracellular bacterial RNA sequencing (<u>Gunnarsson et al., 2025</u>). Briefly, the nf-core dual RNA-seq pipeline (v1.0.0) was modified to allow for UMI deduplication (umi_tools v1.0.1). UMIs from both mates of each read pair were merged and added to corresponding read headers using umi-tools extract. Reads were aligned using STAR to a composite host–pathogen reference genome generated by dual RNA-seq, comprising either the mouse (GRCm39) or human genome (GRCh38, p13) and Mtb H37Rv (NC_000963). The aligned reads were subsequently indexed with samtools, deduplicated using umi-tools dedup, and quantified with HTSeq.

### RNA sequencing analysis

Differential expression was analyzed using DESeq2, and log_2_ fold changes were adjusted using apeGLM (<u>Zhu et al., 2019</u>). Briefly, statistical significance was determined with a Wald test and *p*-values were adjusted using Benjamini-Hochberg correction. Genes with an adjusted *p*-value less than 0.05 and absolute log_2_ fold change greater than 1 were considered significant.

Functional enrichment analysis of Mtb differentially expressed genes was performed using a custom R pipeline implemented in `mtb_enrichment.R` (R version 4.3.3). Functional category annotations were derived from the Mtb H37Rv genome feature (GFF) file downloaded from Mycobrowser (version 5). Genes were assigned to Mycobrowser ‘Functional_Category’ terms by parsing the corresponding attribute field in the GFF file. Genes were defined as differentially expressed (DE) using a false discovery rate (FDR) threshold of 0.05 and absolute log₂FC > 1. Genes with FDR < 0.05 irrespective of fold-change were analyzed in parallel.

For each functional category, enrichment was assessed by a two-sided Fisher’s exact test comparing the proportion of DE genes within the category to that in the genome-wide background. Categories with no annotated genes were excluded. For each term, the odds ratio, *p*-value, and Benjamini–Hochberg–adjusted FDR were calculated. Enrichment direction (enriched or depleted) was assigned based on the relative proportion of DE genes in each category compared to the background.

Results were visualized using a custom ggplot2-based dot panel, in which categories were ranked by FDR and displayed with point size proportional to –log₁₀(FDR) and color denoting enrichment direction. Output tables—including enriched categories, adjusted statistics, and DE gene membership—were exported as Excel workbooks using the openxlsx package. Heatmaps show *z*-scored log-normalized transcripts per million (TPM). Each column represents one host donor. Heatmaps are hierarchically clustered along both axes (by both donors and genes).

## ACKNOWLEDGEMENTS

The biosafety level 3 (BSL-3) work with Mtb strains was performed at the Ragon Institute. The BSL-3 core is supported in part by the Harvard Center for Aids Research (P30 AI060354). We thank Yong Xie, Amy Barczak, and Julie Boucau for directing and managing the BSL-3 facility. We thank Amy Barczak for providing the *eccCa1* transposon mutant and David Sherman for providing the *dosR* mutants. We thank Ryan Milligan for buffy coat processing. We thank all members of the Bryson laboratory for their helpful comments and suggestions during this study. This work was funded by the NIH R01AI166313, R35GM142900 and R01AI184666.

## SUPPLEMENTAL FIGURES CAPTIONS

**Supplemental Figure 1. *M. tuberculosis* forms intracellular lipid inclusions (ILI) during mouse macrophage infection but not in human macrophages. (A)** Human monocyte-derived macrophages (hMDMs) and three murine macrophage types—bone marrow-derived macrophages (BMDMs), immortalized BMDMs (iBMMs), and RAW264.7 macrophages (RAWs)—were infected with cytosolic mCherry-expressing Mtb Erdman for 24 h (MOI = 3 for hMDMs; MOI = 10 for murine macrophages). Following infection, neutral lipids were labeled with BODIPY C16 (8 h pulse, 1 h chase), samples were fixed with paraformaldehyde (PFA), treated with 60% isopropanol to reduce cytoplasmic background fluorescence, and stained with DAPI. Samples were visualized by Airyscan fluorescence microscopy. Scale bars, 10 µm. **(B)** Magnified insets from (A) (iBMMs, gray dashed box) showing nuclei (DAPI, blue), mCherry-labeled Mtb (red), and BODIPY C16-labeled ILIs (green) in separate channels and merged images. Scale bars, 5 µm. **(C)** Quantification of ILI fluorescence from (A), shown as ILI area / Mtb area ± SEM (N = 3). hMDM data are color-coded to indicate independent donors.

**Supplemental Figure 2. *M. tuberculosis* can form ILIs at the different growth stages. (A)** Growth curve of mCherry-expressing Mtb Erdman cultured in 7H9^OADC^ medium, monitored by OD_600_. **(B)** At each day (0–8), cultures were assayed for ILI formation by labeling neutral lipids with BODIPY C16 (4 h pulse, 1 h chase) and visualized by Airyscan fluorescence microscopy. Scale bars, 2 µm. **(C)** Quantification of ILI signal from (B), shown as ILI area / Mtb area ± SEM (N = 3).

**Supplemental Figure 3. Knockout of *M. tuberculosis* ESX-1 secretion system abolishes ILI formation during murine macrophage infection. (A)** Immortalized murine macrophages (iBMMs) were infected with cytosolic mCherry-expressing Mtb H37Rv wild type (wt) or ESX-1 mutant (*eccCa1:Tn*) for 24 h (MOI = 10). After infection, 8 h BODIPY C16 pulse-1h chase experiments were performed. Cells were fixed, stained with DAPI, and visualized using Airyscan fluorescence microscopy. Scale bars, 10 µm. **(B)** Quantification of ILI signal from (A), presented as ILI area / Mtb area ± SEM (N = 3). **(C)** mCherry-expressing Mtb H37Rv wt and *eccCa1:Tn* strains were cultured axenically in 7H9^OADC^ medium and labeled with BODIPY C16 (4 h pulse, 1 h chase) at OD₆₀₀ ≈ 1. Samples were fixed and visualized by Airyscan fluorescence microscopy. Scale bars, 5 µm. All data represent at least three independent experiments.

## REFERENCES

Ahmed, M., Thirunavukkarasu, S., Rosa, B.A., Thomas, K.A., Das, S., Rangel-Moreno, J., Lu, L., Mehra, S., Mbandi, S.K., Thackray, L.B., et al. (2020). Immune correlates of tuberculosis disease and risk translate across species. Sci Transl Med 12. **doi:**10.1126/scitranslmed.aay0233

Batt, S.M., Abrahams, K.A., & Besra, G.S. (2024). Top five unanswered questions in bacterial cell wall research. Cell Surf 11, 100122. **doi:**10.1016/j.tcsw.2024.100122

Borah, K., Mendum, T.A., Hawkins, N.D., Ward, J.L., Beale, M.H., Larrouy-Maumus, G., Bhatt, A., Moulin, M., Haertlein, M., Strohmeier, G., et al. (2021). Metabolic fluxes for nutritional flexibility of *Mycobacterium tuberculosis*. Mol Syst Biol 17, e10280. **doi:**10.15252/msb.202110280

Chen, F., Lukat, P., Iqbal, A.A., Saile, K., Kaever, V., van den Heuvel, J., Blankenfeldt, W., Bussow, K., & Pessler, F. (2019). Crystal structure of *cis*-aconitate decarboxylase reveals the impact of naturally occurring human mutations on itaconate synthesis. Proc Natl Acad Sci U S A 116, 20644–20654. **doi:**10.1073/pnas.1908770116

Comas, I., Coscolla, M., Luo, T., Borrell, S., Holt, K.E., Kato-Maeda, M., Parkhill, J., Malla, B., Berg, S., Thwaites, G., et al. (2013). Out-of-Africa migration and Neolithic coexpansion of *Mycobacterium tuberculosis* with modern humans. Nat Genet 45, 1176–1182. **doi:**10.1038/ng.2744

Cooper, A.M., Pearl, J.E., Brooks, J.V., Ehlers, S., & Orme, I.M. (2000). Expression of the nitric oxide synthase 2 gene is not essential for early control of *Mycobacterium tuberculosis* in the murine lung. Infect Immun 68, 6879–6882. **doi:**10.1128/IAI.68.12.6879-6882.2000

Daniel, J., Maamar, H., Deb, C., Sirakova, T.D., & Kolattukudy, P.E. (2011). *Mycobacterium tuberculosis* uses host triacylglycerol to accumulate lipid droplets and acquires a dormancy-like phenotype in lipid-loaded macrophages. PLoS Pathog 7, e1002093. **doi:**10.1371/journal.ppat.1002093

Gago, G., Diacovich, L., & Gramajo, H. (2018). Lipid metabolism and its implication in mycobacteria-host interaction. Curr Opin Microbiol 41, 36–42. **doi:**10.1016/j.mib.2017.11.020

Garton, N.J., Christensen, H., Minnikin, D.E., Adegbola, R.A., & Barer, M.R. (2002). Intracellular lipophilic inclusions of mycobacteria *in vitro* and in sputum. Microbiology (Reading) 148, 2951–2958. **doi:**10.1099/00221287-148-10-2951

Gilbertson, S.E., Walter, H.C., Gardner, K., Wren, S.N., Vahedi, G., & Weinmann, A.S. (2022). Topologically associating domains are disrupted by evolutionary genome rearrangements forming species-specific enhancer connections in mice and humans. Cell Rep 39, 110769. **doi:**10.1016/j.celrep.2022.110769

Greenwood, D.J., Dos Santos, M.S., Huang, S., Russell, M.R.G., Collinson, L.M., MacRae, J.I., West, A., Jiang, H., & Gutierrez, M.G. (2019). Subcellular antibiotic visualization reveals a dynamic drug reservoir in infected macrophages. Science 364, 1279–1282. **doi:**10.1126/science.aat9689

Groschel, M.I., Sayes, F., Simeone, R., Majlessi, L., & Brosch, R. (2016). ESX secretion systems: mycobacterial evolution to counter host immunity. Nat Rev Microbiol 14, 677–691. **doi:**10.1038/nrmicro.2016.131

Gunnarsson, C., McGinn, R.A., Hochfelder, J., & Bryson, B.D. (2025). Systems-level characterization of EGFR kinase inhibitors reveals heterogeneous effects on Mtb-macrophage interactions. bioRxiv preprint. **doi:**10.1101/2025.10.17.683041

Knight, M., Braverman, J., Asfaha, K., Gronert, K., & Stanley, S. (2018). Lipid droplet formation in *Mycobacterium tuberculosis* infected macrophages requires IFN-gamma/HIF-1alpha signaling and supports host defense. PLoS Pathog 14, e1006874. **doi:**10.1371/journal.ppat.1006874

Lee, W., VanderVen, B.C., Fahey, R.J., & Russell, D.G. (2013). Intracellular *Mycobacterium tuberculosis* exploits host-derived fatty acids to limit metabolic stress. Journal of Biological Chemistry 288, 6788–6800. **doi:**10.1074/jbc.M112.445056

Luthra, A., Mahmood, A., Arora, A., & Ramachandran, R. (2008). Characterization of *Rv3868*, an essential hypothetical protein of the ESX-1 secretion system in *Mycobacterium tuberculosis*. Journal of Biological Chemistry 283, 36532–36541. **doi:**10.1074/jbc.M807144200

Mallick, I., Santucci, P., Poncin, I., Point, V., Kremer, L., Cavalier, J.F., & Canaan, S. (2021). Intrabacterial lipid inclusions in mycobacteria: unexpected key players in survival and pathogenesis? FEMS Microbiol Rev 45. **doi:**10.1093/femsre/fuab029

Mehra, S., Foreman, T.W., Didier, P.J., Ahsan, M.H., Hudock, T.A., Kissee, R., Golden, N.A., Gautam, U.S., Johnson, A.M., Alvarez, X., et al. (2015). The DosR Regulon Modulates Adaptive Immunity and Is Essential for *Mycobacterium tuberculosis* Persistence. Am J Respir Crit Care Med 191, 1185–1196. **doi:**10.1164/rccm.201408-1502OC

Michelucci, A., Cordes, T., Ghelfi, J., Pailot, A., Reiling, N., Goldmann, O., Binz, T., Wegner, A., Tallam, A., Rausell, A., et al. (2013). Immune-responsive gene 1 protein links metabolism to immunity by catalyzing itaconic acid production. Proc Natl Acad Sci U S A 110, 7820–7825. **doi:**10.1073/pnas.1218599110

Mills, E.L., Ryan, D.G., Prag, H.A., Dikovskaya, D., Menon, D., Zaslona, Z., Jedrychowski, M.P., Costa, A.S.H., Higgins, M., Hams, E., et al. (2018). Itaconate is an anti-inflammatory metabolite that activates Nrf2 via alkylation of KEAP1. Nature 556, 113–117. **doi:**10.1038/nature25986

Nair, S., Huynh, J.P., Lampropoulou, V., Loginicheva, E., Esaulova, E., Gounder, A.P., Boon, A.C.M., Schwarzkopf, E.A., Bradstreet, T.R., Edelson, B.T., et al. (2018). *Irg1* expression in myeloid cells prevents immunopathology during *M. tuberculosis* infection. J Exp Med 215, 1035–1045. **doi:**10.1084/jem.20180118

Nazarova, E.V., Podinovskaia, M., Russell, D.G., & VanderVen, B.C. (2018). Flow Cytometric Quantification of Fatty Acid Uptake by *Mycobacterium tuberculosis* in Macrophages. Bio Protoc 8. **doi:**10.21769/bioprotoc.2734

Ohno, H., Zhu, G., Mohan, V.P., Chu, D., Kohno, S., Jacobs, W.R., Jr., & Chan, J. (2003). The effects of reactive nitrogen intermediates on gene expression in *Mycobacterium tuberculosis*. Cell Microbiol 5, 637–648. **doi:**10.1046/j.1462-5822.2003.00307.x

Pandey, A.K., & Sassetti, C.M. (2008). Mycobacterial persistence requires the utilization of host cholesterol. Proc Natl Acad Sci U S A 105, 4376–4380. **doi:**10.1073/pnas.0711159105

Park, H.D., Guinn, K.M., Harrell, M.I., Liao, R., Voskuil, M.I., Tompa, M., Schoolnik, G.K., & Sherman, D.R. (2003). *Rv3133c*/*dosR* is a transcription factor that mediates the hypoxic response of *Mycobacterium tuberculosis*. Molecular Microbiology 48, 833–843. **doi:**10.1046/j.1365-2958.2003.03474.x

Peyron, P., Vaubourgeix, J., Poquet, Y., Levillain, F., Botanch, C., Bardou, F., Daffe, M., Emile, J.F., Marchou, B., Cardona, P.J., et al. (2008). Foamy macrophages from tuberculous patients’ granulomas constitute a nutrient-rich reservoir for *M. tuberculosis* persistence. PLoS Pathog 4, e1000204. **doi:**10.1371/journal.ppat.1000204

Pisu, D., Huang, L., Rin Lee, B.N., Grenier, J.K., & Russell, D.G. (2020). Dual RNA-Sequencing of *Mycobacterium tuberculosis*-Infected Cells from a Murine Infection Model. STAR Protoc 1, 100123. **doi:**10.1016/j.xpro.2020.100123

Podinovskaia, M., Lee, W., Caldwell, S., & Russell, D.G. (2013). Infection of macrophages with *Mycobacterium tuberculosis* induces global modifications to phagosomal function. Cell Microbiol 15, 843–859. **doi:**10.1111/cmi.12092

Priya, M., Gupta, S.K., Koundal, A., Kapoor, S., Tiwari, S., Kidwai, S., Sorio de Carvalho, L.P., Thakur, K.G., Mahajan, D., Sharma, D., et al. (2025). Itaconate mechanism of action and dissimilation in *Mycobacterium tuberculosis*. Proc Natl Acad Sci U S A 122, e2423114122. **doi:**10.1073/pnas.2423114122

Russell, D.G., Cardona, P.J., Kim, M.J., Allain, S., & Altare, F. (2009). Foamy macrophages and the progression of the human tuberculosis granuloma. Nat Immunol 10, 943–948. **doi:**10.1038/ni.1781

Schnappinger, D., Ehrt, S., Voskuil, M.I., Liu, Y., Mangan, J.A., Monahan, I.M., Dolganov, G., Efron, B., Butcher, P.D., Nathan, C., et al. (2003). Transcriptional Adaptation of *Mycobacterium tuberculosis* within Macrophages: Insights into the Phagosomal Environment. J Exp Med 198, 693–704. **doi:**10.1084/jem.20030846

Schroder, K., Irvine, K.M., Taylor, M.S., Bokil, N.J., Le Cao, K.A., Masterman, K.A., Labzin, L.I., Semple, C.A., Kapetanovic, R., Fairbairn, L., et al. (2012). Conservation and divergence in Toll-like receptor 4-regulated gene expression in primary human versus mouse macrophages. Proc Natl Acad Sci U S A 109, E944–953. **doi:**10.1073/pnas.1110156109

Shi, X., Zhou, H., Wei, J., Mo, W., Li, Q., & Lv, X. (2022). The signaling pathways and therapeutic potential of itaconate to alleviate inflammation and oxidative stress in inflammatory diseases. Redox Biol 58, 102553. **doi:**10.1016/j.redox.2022.102553

Thomas, S.T., Wierenga, K.A., Pestka, J.J., & Olive, A.J. (2022). Fetal Liver-Derived Alveolar-like Macrophages: A Self-Replicating *Ex Vivo* Model of Alveolar Macrophages for Functional Genetic Studies. Immunohorizons 6, 156–169. **doi:**10.4049/immunohorizons.2200011

Voskuil, M.I., Schnappinger, D., Visconti, K.C., Harrell, M.I., Dolganov, G.M., Sherman, D.R., & Schoolnik, G.K. (2003). Inhibition of respiration by nitric oxide induces a *Mycobacterium tuberculosis* dormancy program. J Exp Med 198, 705–713. **doi:**10.1084/jem.20030205

Waltermann, M., Hinz, A., Robenek, H., Troyer, D., Reichelt, R., Malkus, U., Galla, H.J., Kalscheuer, R., Stoveken, T., von Landenberg, P., et al. (2005). Mechanism of lipid-body formation in prokaryotes: how bacteria fatten up. Molecular Microbiology 55, 750–763. **doi:**10.1111/j.1365-2958.2004.04441.x

Waltermann, M., & Steinbuchel, A. (2005). Neutral lipid bodies in prokaryotes: recent insights into structure, formation, and relationship to eukaryotic lipid depots. Journal of Bacteriology 187, 3607–3619. **doi:**10.1128/JB.187.11.3607-3619.2005

Warner, D.F., Barczak, A.K., Gutierrez, M.G., & Mizrahi, V. (2025). *Mycobacterium tuberculosis* biology, pathogenicity and interaction with the host. Nat Rev Microbiol. **doi:**10.1038/s41579-025-01201-x

Waters, J.K., & Eijkelkamp, B.A. (2024). Bacterial acquisition of host fatty acids has far-reaching implications on virulence. Microbiol Mol Biol Rev 88, e0012624. **doi:**10.1128/mmbr.00126-24

Wirth, T., Hildebrand, F., Allix-Beguec, C., Wolbeling, F., Kubica, T., Kremer, K., van Soolingen, D., Rusch-Gerdes, S., Locht, C., Brisse, S., et al. (2008). Origin, spread and demography of the *Mycobacterium tuberculosis* complex. PLoS Pathog 4, e1000160. **doi:**10.1371/journal.ppat.1000160

Yen, C.L., Stone, S.J., Koliwad, S., Harris, C., & Farese, R.V., Jr. (2008). Thematic review series: glycerolipids. DGAT enzymes and triacylglycerol biosynthesis. J Lipid Res 49, 2283–2301. **doi:**10.1194/jlr.R800018-JLR200

Zhang, Y.J., Reddy, M.C., Ioerger, T.R., Rothchild, A.C., Dartois, V., Schuster, B.M., Trauner, A., Wallis, D., Galaviz, S., Huttenhower, C., et al. (2013). Tryptophan biosynthesis protects mycobacteria from CD4 T-cell-mediated killing. Cell 155, 1296–1308. **doi:**10.1016/j.cell.2013.10.045

Zhu, A., Ibrahim, J.G., & Love, M.I. (2019). Heavy-tailed prior distributions for sequence count data: removing the noise and preserving large differences. Bioinformatics 35, 2084–2092. **doi:**10.1093/bioinformatics/bty895

