## Supplementary figures and images for "Systems analysis reveals alternate metabolic states adopted by *Mycobacterium tuberculosis* across species"

### Supplemental Figure 1

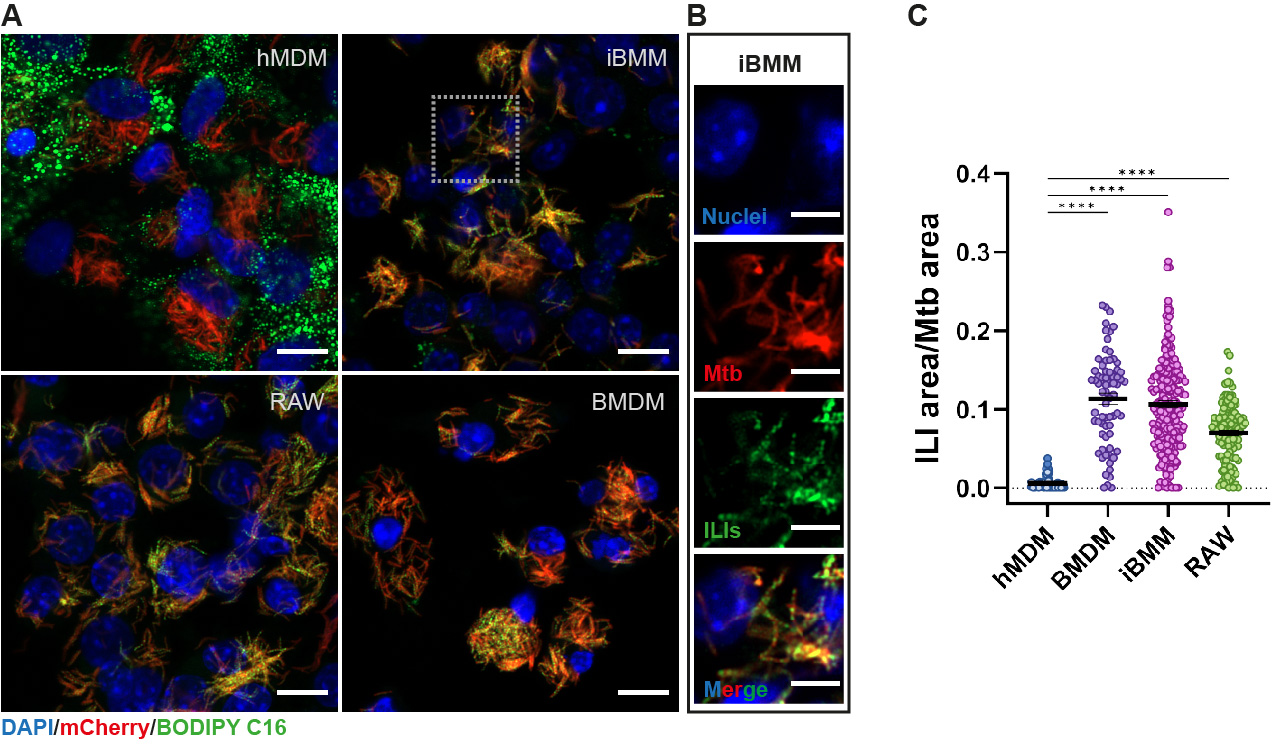

### Supplemental Figure 2

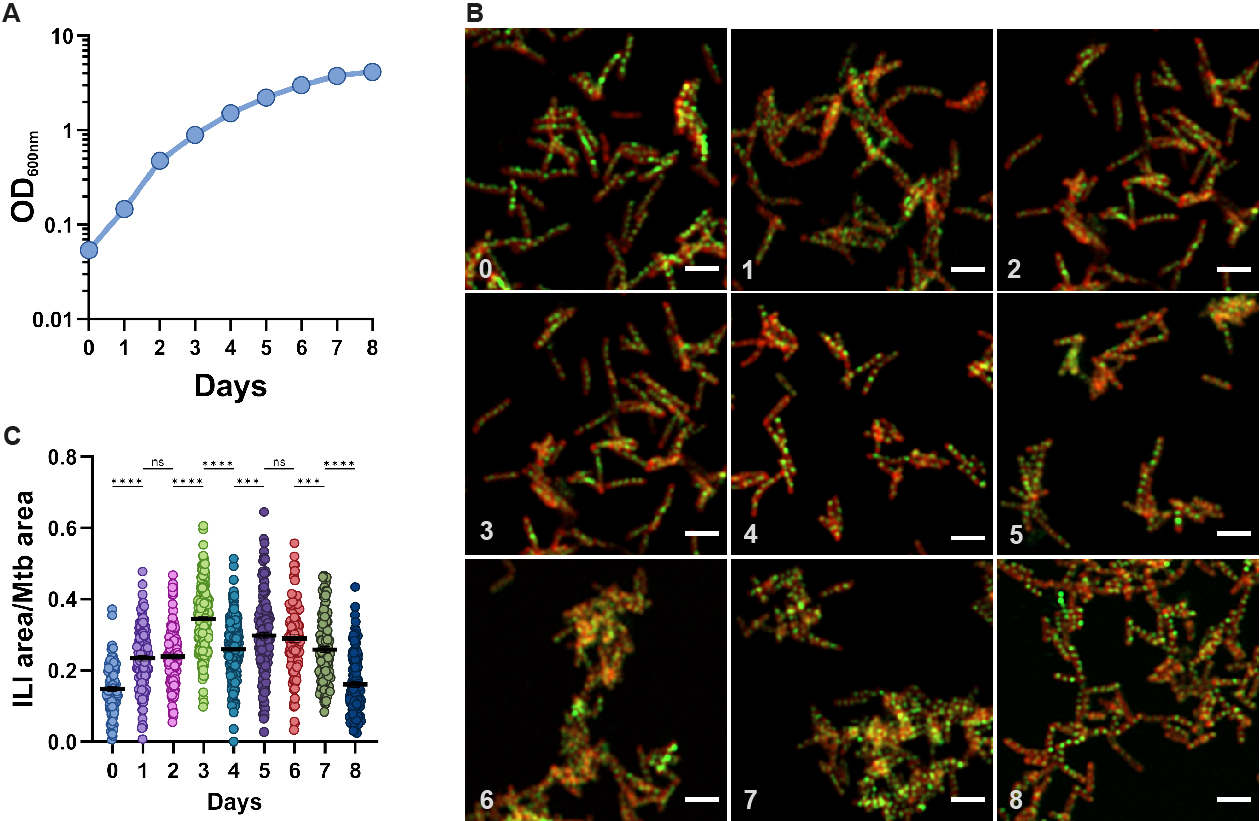

### Supplemental Figure 3

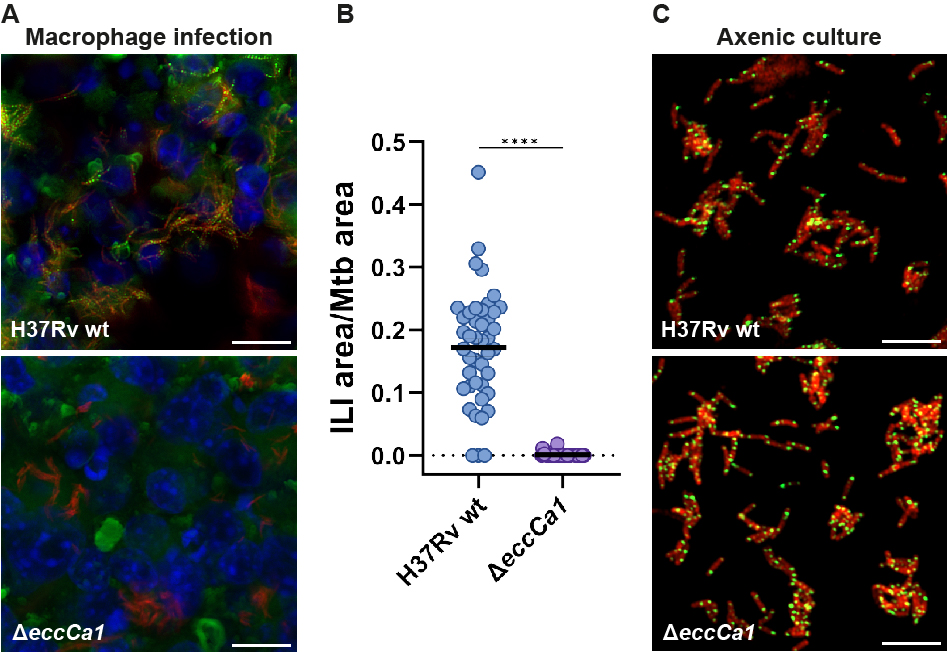
